# Dynamic medial parietal and hippocampal deactivations under DMT relate to sympathetic output and altered sense of time, space, and the self

**DOI:** 10.1101/2024.02.14.580356

**Authors:** Lorenzo Pasquini, Alexander J. Simon, Courtney L. Gallen, Hannes Kettner, Leor Roseman, Adam Gazzaley, Robin L. Carhart-Harris, Christopher Timmermann

## Abstract

N,N-Dimethyltryptamine (DMT) is a serotonergic psychedelic, known to rapidly induce short-lasting alterations in conscious experience, characterized by a profound and immersive sense of physical transcendence alongside rich and vivid auditory distortions and visual imagery. Multimodal neuroimaging data paired with dynamic analysis techniques offer a valuable approach for identifying unique signatures of brain activity – and linked autonomic physiology – naturally unfolding during the altered state of consciousness induced by DMT. We leveraged simultaneous fMRI and EKG data acquired in 14 healthy volunteers prior to, during, and after intravenous administration of DMT, and, separately, placebo. fMRI data was preprocessed to derive individual dynamic activity matrices, reflecting the similarity of brain activity in time, and community detection algorithms were applied on these matrices to identify brain activity substates; EKG data was used to derive continuous heart rate. We identified a brain substate occurring immediately after DMT injection, characterized by hippocampal and medial parietal deactivations and increased superior temporal lobe activity under DMT. Deactivations in the hippocampus and medial parietal cortex correlated with alterations in the usual sense of time, space and self-referential processes, reflecting a deconstruction of essential features of ordinary consciousness. Superior lobe activations instead correlated with audio/visual hallucinations and experience of “*entities*”, reflecting the emergence of altered sensory experiences under DMT. Finally, increased heart rate under DMT correlated positively with hippocampus/medial parietal deactivation and the experience of “*entities*”, and negatively with altered self-referential processes. These results suggest a chain of influence linking sympathetic regulation to hippocampal and medial parietal deactivations under DMT, which combined, may contribute to positive mental health outcomes related to self-referential processing following psychedelic administration.

## Introduction

Bodily self-awareness (Damasio and Damasio, 2022) is thought to result from the integration of visceral and autonomic bodily functions into a distributed brain network supporting homeostasis, interoception, and emotions (Critchley and Harrison, 2013), encompassing a set of subcortical and cortical structures including the insula, anterior cingulate cortex, and ventral striatum (Seeley et al., 2007). Conversely, one’s sense of identity rooted in autobiographic memories, often referred to as the “narrative self” (Gallagher, 2000), has been demonstrated to rely on a distributed brain network spanning midline parietal and medial temporal structures, including the precuneus, posterior cingulate cortex, and hippocampi (Northoff and Bermpohl, 2004). These regions are collectively known to be a core component of the default mode network (DMN) (Raichle et al., 2001), a large-scale brain system associated with self-referential processes such as autobiographical memories, mind wandering, and future planning (Buckner and DiNicola, 2019). Crucially, the investigation of the neural underpinning of one’s sense of self has been catalyzed via the timely re-introduction of psychedelic substances in human neuroscience research (Vollenweider and Kometer, 2010; Carhart-Harris et al., 2012; McCulloch et al., 2022).

N,N-Dimethyltryptamine (DMT) is a serotonergic psychedelic (Nichols, 2016) known to rapidly induce an intense but short-lasting altered state of consciousness (Strassman, 1995; Vogt et al., 2023), characterized by a sense of transcendence of physical bounds and intense sensory immersion – including vivid and elaborate visual imagery, and in about half of cases, a sense of being in the presence of other sentient entities (Lawrence et al., 2022; Timmermann et al., 2023b). The subjective effects of DMT, when administered intravenously at high doses, arise within seconds of the injection, and rapidly progress into an altered state of consciousness characterized by deep and profound immersion (Timmermann et al., 2019). This state typically lasts several minutes, and gradually attenuates as participants regain normal waking consciousness approximately 20 min following the injection (Timmermann et al., 2023b). Due to its fast dynamics and profound subjective effects, DMT has been recognized as a powerful tool to explore the neurophenomenology and neural underpinning of consciousness (Timmermann et al., 2023a). as it deconstructs essential features of conscious experience (such as the sense of time, space and self) and catalyzes the emergence of novel content (such as visual imagery and the experience of entities).

As with other serotonergic psychedelics, the altered state of consciousness induced by DMT is usually accompanied by changes in autonomic and central nervous system physiology (Strassman and Qualls, 1994; Carbonaro and Gatch, 2016; Alamia et al., 2020). DMT often triggers transient increases in sympathetic tone as measured through continuous heart rate, which normalize as the effects of the drug dissipate (Strassman and Qualls, 1994). Further, recent studies have found associations between altered cardiac activity and the psychedelic experience (Rosas et al., 2023). Studies leveraging resting-state functional magnetic resonance imaging (rs-fMRI), a method used to measure the synchronicity of spontaneous activity across distant brain regions (Smith et al., 2009), have shown that DMT drives global hyperconnectivity, collapses hierarchical organization, and reduces intra-network integrity, particularly among core regions of the DMN (Timmermann et al., 2023b). Further, dynamic analyses have revealed that rs-fMRI-based global connectivity peaks within the first 4-6 minutes after the injection, corresponding to the highest intensity ratings of subjective effects under DMT (Timmermann et al., 2023b). These findings highlight the dynamic character of DMT and raise further questions about whether distinguishable physiological substates could be identified across the natural progression of a psychedelic experience under DMT. Novel dynamic analysis methods allow for the identification of functional substates occurring during the duration of an entire scan (Deco et al., 2008; Calhoun et al., 2014; Lurie et al., 2020). Dynamic analysis approaches could hence play an invaluable contribution to the identification of distinguishable signatures of brain activity – and linked autonomic physiology – naturally emerging and dissolving under the fast action of DMT.

Here, we applied dynamic analysis methods combined with graph theoretical techniques to rs-fMRI and electrocardiogram (EKG) data simultaneously acquired in a pharmacological study with 14 healthy volunteers, to assess the natural evolution of autonomic physiology and brain activity substates during the administration of DMT.

## Materials and Methods

### Study sample

This study involved secondary analyses of fMRI and EKG data simultaneously acquired in 14 healthy volunteers (4 females; age [mean, SD, range] in years = 34.1, 8.8, 23-53) in the context of a single-blind, placebo-controlled, counter-balanced study assessing the effects of DMT on brain function (Timmermann et al., 2023b). Primary exclusion criteria were: <18 years of age, MRI contraindications, absence of experience with a psychedelic, an adverse reaction to a psychedelic, history of psychiatric or physical illness rendering unsuitable for participation (i.e., diabetes, epilepsy, or heart disease), family history of psychotic disorder, or excessive use of alcohol or drugs of abuse. All participants provided written informed consent for participation in the study. This study was approved by the National Research Ethics Committee London — Brent and the Health Research Authority and was conducted under the guidelines of the revised Declaration of Helsinki (2000), the International Committee on Harmonization Good Clinical Practices guidelines, and the National Health Service Research Governance Framework. Imperial College London sponsored the research, which was conducted under a Home Office license for research with Schedule 1 drugs.

### Study design

20 enrolled participants visited the Imperial College Clinical Imaging Facility and completed two visits separated by two weeks. At each visit, participants were tested for drugs of abuse and completed two separate sessions: a “resting state” and a “ratings” session. For the “resting state” session, participants were placed in an MRI scanner after placing MRI-compatible EKG sensors and an electroencephalography (EEG) cap was placed on their scalp. The acquired EEG data was analyzed in previous work, and we did not further analyze this data since it was outside the scope of the current study. While being scanned, participants received intravenous administration of either placebo (10 mL of sterile saline) or 20 mg DMT (in fumarate form dissolved in 10 mL of sterile saline) — injected over 30 s, and then flushed with 10 mL of saline over 15 s — in a counter-balanced order (half of the participants received placebo and the other half received DMT for this session). Rs-fMRI data acquisition lasted 28 min in total, with DMT/placebo being administered at the end of the 8th min. While participants laid in the scanner with their eyes closed, EKG activity was simultaneously recorded through MRI compatible sensors. At the end of the scanning session, participants were interviewed and completed questionnaires designed to assess the subjective effects experienced during the scan. A second session then followed with the same procedure as the initial session, except on this occasion, participants were (audio) cued to verbally rate the subjective intensity of drug effects every minute in real time while in the scanner. A second visit consisted of the same procedure as the first visit, however the order of DMT/placebo administration was reversed for each of the sessions. This article reports the results concerning the fMRI/EKG data collected during resting-state scans where participants received either DMT or a placebo and were not interrupted for experience sampling purposes. Only continuously assessed intensity ratings were used in this study to show when identified brain activity substates occurred in relationship to the strength of the experience under DMT. Overall, the MRI environment was well tolerated, and dosing events were not accompanied by adverse events (see also ***Supplementary Methods*** for details regarding participant safety)(Petitmengin, 2006; Johnson et al., 2008).

### Visual analogue scales rating at the end of the scanning session

After participants completed the scanning session and regained normal waking consciousness, they were asked to rate several items describing the DMT/placebo experience using visual analogue scales (ratings ranged from 0-1, in incremental steps of 0.01). 25 items were presented sequentially, which assessed aspects of the psychedelic experience related to drug intensity, sensory distortion (including visual and auditory hallucinations), changes in time and space, feelings of disembodiment and dissociation, and self-referential processes (e.g., freely wandering thoughts, ego-dissolution, and meaningfulness of the experience) (Timmermann et al., 2023b). Ratings for theses scales were acquired for both the DMT and the placebo conditions. The 11 Dimensions Altered States of Consciousness Questionnaire (Studerus et al., 2010) and the 30-item Mystical Experience Questionnaire (Barrett et al., 2015) were also assessed over the course of the day, but where not analyzed in the context of the current study, since we focused on the visual analogue scales acquired immediately after the end of the scanning session.

### EKG

EKG data was leveraged to derive continuous heart rate estimates. EKG data was collected simultaneously to the fMRI scan using two electrodes, one placed on the participant’s back (behind the chest area), and the other placed above the heart area. The data was recorded with an MR-compatible BrainAmp MR amplifier (BrainProducts GmbH, Munich, Germany) using a 5 kHz sampling rate and a 250 Hz low-pass filter. EKG-MRI clock synchronization was ensured using the Brain Products SyncBox hardware. As for fMRI, recordings lasted 28 min, with 8 min of baseline and 20 min post injection (see Timmermann et al., 2023b for details).

EKG data acquired simultaneously to the MRI was preprocessed using custom code in MATLAB (https://www.mathworks.com/products/matlab.html)(Supplementary ***Figure S1***). After visually inspecting the raw data, inter-beat intervals were estimated from the EKG via a semi-automated procedure where first negative voltages were set to 0, to improve the detection of R peaks. The positive voltage EKG was then processed via a wavelet transform, and then peaks where detected using the *findpeaks* function in MATLAB with manually adjusted minimum height (7,500 microvolts) and minimum inter-beat distance (0.45 seconds) (Goldberger et al., 2000; Moody and Mark, 2001; Rosas et al., 2023). Subsequently, continuous heart rate was estimated in beats per minute (bpm) as follows:

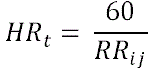

where *HR_t_* is the instantaneous heart rate, 60 reflects the average heart rate in bpm, and *RR_ij_* reflects the interval in seconds between subsequent R-peaks *i* and *j* (Pasquini et al., 2022). Continuous heart-rate estimates were down-sampled from 250 Hz to 0.5 Hz, the same sampling frequency as fMRI data.

### Neuroimaging

#### Neuroimaging data acquisition

Functional images were acquired in a 3T MRI scanner (Siemens Magnetom Verio syngo MR B17) using a 12-channel head coil compatible with EEG acquisition using a T2*-weighted blood-oxygen-level-dependent (BOLD) sensitive gradient echo planar imaging sequence [repetition time (TR) = 2000ms, echo time (TE) = 30 ms, acquisition time (TA) = 28.06 mins, flip angle (FA) = 80°, voxel size = 3.0 × 3.0 × 3.0mm3, 35 slices, interslice distance = 0 mm]. Whole-brain T1-weighted structural images were also acquired.

#### Neuroimaging data preprocessing

Details on MRI data preprocessing can be found on previous published work (Carhart-Harris et al., 2016) and in the parent paper (Timmermann et al., 2023b). Briefly, preprocessing leveraged standard AFNI (https://afni.nimh.nih.gov/), FSL (https://fsl.fmrib.ox.ac.uk/fsl/fslwiki), and in house code to perform rs-fMRI data despiking, slice time correction, functional registration for motion correction, anatomical brain extraction, rigid body registration of functional scans to anatomical scans, and nonlinear registration to a 2 mm Montreal Neurological Institute brain template. Additional preprocessing steps included rs-fMRI data scrubbing using a framewise head-displacement threshold of 0.4 mm followed by linear interpolation using the mean of the surrounding volumes, spatial smoothing with a 6 mm full width at half maximum kernel, band-pass filtering between 0.01 and 0.08 Hz, linear and quadratic detrending, and regression of nine motion-related and three anatomical nuisance regressors (Timmermann et al., 2023b). Participants with >20% of scrubbed volumes based on a frame-wise displacement threshold of 0.4 mm were discarded from group analyses, resulting in the final sample of 14 participants analyzed in this study, of which 7 received DMT at the first visit and placebo at the second visit, while the remaining 7 received placebo at the first visit and DMT at the second visit.

#### Neuroimaging data analysis

Voxel-level brain activity estimates derived from rs-fMRI data were reduced by obtaining the average activity from 100 areas in the Schaefer atlas (Schaefer et al., 2018) and 12 subcortical areas derived from the automated anatomical atlas (AAL) (Tzourio-Mazoyer et al., 2002). Pearson’s correlation was used to derive individual time-resolved brain activity similarity matrices, by, e.g., correlating the activity of all nodes at time point one with the activity of all nodes at time point two. This procedure resulted in matrices reflecting the homogeneity of brain activity during the entire duration of the scanning session (Kringelbach and Deco, 2020), estimated once for the DMT and once for the placebo conditions (**Figure 1A-B**). Group-averaged time-resolved brain activity similarity matrices derived once under DMT and once under placebo were then subtracted, and these subtraction matrices were used to identify brain activity substates differentiating DMT from placebo using a fully data driven approach (**Figure 1C**) as described in the following section.

**Figure 1.**
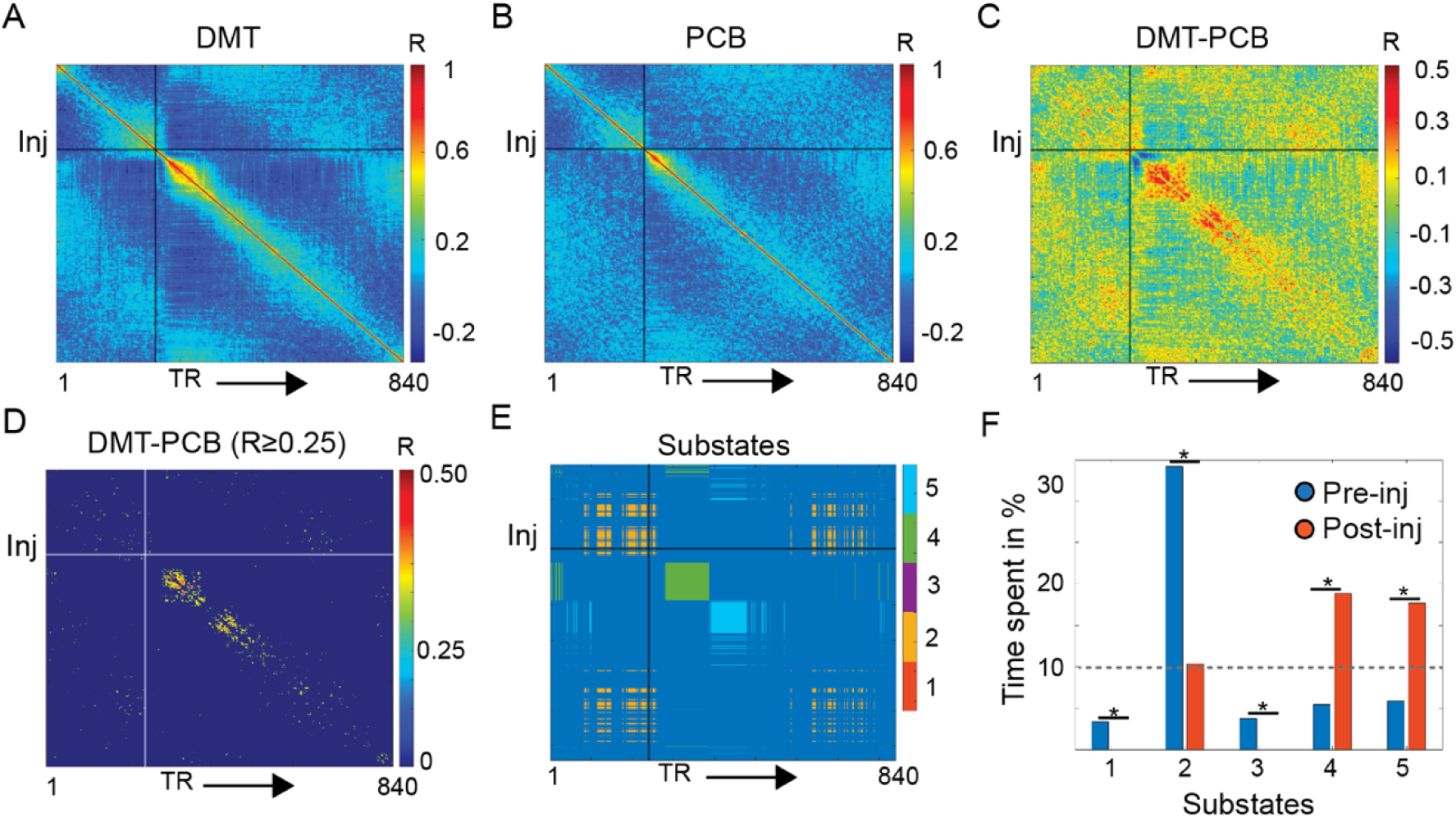
Analysis pipeline and metrics. Group-mean continuous brain activity similarity matrix, reflecting the homogeneity of brain activity in time during the DMT (**A**) and the placebo (PCB; **B**) conditions. (**C**) Mean subtraction matrix reflecting how continuous brain activity similarity varies across the DMT and placebo conditions. Warm colors reflect higher similarity of brain activity during the DMT condition. (**D**) The subtraction matrix was thresholded for similarity values maximizing the identification of separate brain activity substates occurring either during the pre-injection or post-injection periods (Pearson’s correlation value R ≥ 0.25) using an unsupervised community detection algorithm. This community detection algorithm identified five brain activity substates differentiating the DMT from the placebo condition (**E**). (**F**) These five substates were occupied at significant higher rates either during the pre-or post-injection periods, but only three out of these five substates were occupied for more that 10% of the duration of the pre-or post-injection scanning time. Following analyses focused on these three activation substates: State 2, 4, and 5. DMT = N,N-Dimethyltryptamine; Inj = timepoint of injection, indicated by continuous vertical and horizontal lines; TR = fMRI volume acquired at each repetition time. \**p* < 0.05

The subtraction matrix was iteratively thresholded for absolute Pearson’s correlation values ranging from 0.20 to 0.30, in incremental steps of 0.01 (**Figure 1D** and ***Supplementary Figure S2A-C***). A range between 0.20-0.30 was chosen based on the right-tailed distribution of these values (***Supplementary Figure S2A***). For each of these thresholded matrices, the Louvain community detection algorithm, as implemented in the publicly available Brain Connectivity Toolbox (https://sites.google.com/site/bctnet/)(Rubinov and Sporns, 2010), was then used to identify brain activity substates within the duration of the entire scan (**Figure 1E**). Community detection was implemented using the default resolution parameter *G* = 1 and an asymmetric treatment of negative weights. This procedure resulted in community affiliation vectors, assigning each functional volume to one of the identified brain activity substates. The community affiliation vectors were then used to separately quantify the occupancy of brain activity substates before and after the injection. This was attained by dividing the occurrence of a specific brain activity substate before/after the injection by the amount of rs-fMRI scans before/after the injection. *X^2^* tests, quantifying which brain activity substates were significantly more occupied either during the pre-or the post-injection periods (*p* < 0.05), were used to identify a subset of brain activity substates that were sufficiently occupied, as well as differentially occupied, when comparing the pre-to the post-injection periods (**Figure 1F** and ***Supplementary Figure S2A-C***). The distinct thresholds applied on the subtraction matrix yielded a varying number of brain activity substates for each iteration (between 2-6), which were differentially occupied prior and after the injection (as quantified through *X*^2^ tests). An R threshold of 0.25 was chosen since it maximized the number of identified brain activity substates substantially occupying either the pre-or post-injection periods (***Supplementary Figure S2B***). For control analyses, brain activity substates were also investigated using thresholds of 0.24 and 0.26, which also yielded high numbers of substates differentially occupying the pre– and post-injection periods (***Supplementary Figure S2C***).

The community affiliation vectors resulting from applying the community detection algorithm on the subtraction matrix thresholded with the optimal solution where then used to derive mean brain activity maps for each participant. These community affiliation vectors were also entered in a general linear model, using individual nodal activity time series as dependent variables, to derive nodal β-maps reflecting individual estimates of brain activity substates (***Supplementary Figure S3***).

### Statistical Analyses

Individual nodal β-maps of brain activity substates were compared across the placebo and DMT conditions using paired *t*-tests (false discovery rate [FDR] adjusted *p* < 0.05). This procedure helped identify a brain activity substate, which emerged immediately after DMT injection and lasted for several minutes, that showed significant differences in nodal activity across the DMT and placebo conditions. Nodes of this brain activity substate displaying either hyperactivity or hypoactivity in the DMT condition were identified and then used as mask to derive averaged levels of activity from the individual β-maps. Identified nodes were also used on the preprocessed rs-fMRI volumes to estimate averaged BOLD activity time series of these regions for the duration of the entire rs-fMRI scan. Individual estimates of nodal hyperactivity or hypoactivity under DMT were entered as dependent variables in a partial least-square regression analysis, using questionnaire change scores (DMT minus placebo) as the set of predicting variables. This method was used due to partial least-squares regression being ideally suited to reduce large numbers of variables, used to predict, to a smaller set of predictors. The partial least-square regression was conducted by estimating two components using the *plsregress* function in MATLAB. Ratings of subjective experience scales were compared across the DMT and placebo conditions using paired *t*-tests (*p* < 0.05 FDR corrected for multiple comparisons). Individual heart rate was averaged during the occurrence of the previously mentioned post-injection brain activity substate and compared across the placebo and DMT conditions using a paired *t*-test (*p* < 0.05). Changes in heart rate during that substate were associated to nodal hypoactivity changes using Pearson’s correlation coefficients (*p* < 0.05). A separate partial least-square regression analysis was used to assess the link between heart rate changes during the first post-injection substate (dependent variable) and questionnaire change scores (predicting variables). As in the previous analysis, two components were estimated. Pearson’s correlation (*p* < 0.05) was used to test the link between absolute mean heart rate changes during the first post-injection substate and mean heart rate during that substate on placebo. A median split was finally used to separate participants into individuals showing low heart rate increase or high heart rate increase during specific brain activity substates emerging under DMT. Three linear regression models were used to test the potential link between order of DMT dosing (whether at the first or at the second visit) and change in brain activity and mean heart rate during the first post-injection substate.

Given the small samples size of the study, a post-hoc power analysis was performed using the publicly available toolbox G*Power v3.1 (Faul et al., 2007) (***Supplementary Analyses***).

## Results

### Dynamic brain activity substates under DMT and placebo

In this study, we explored the dynamic emergence of brain activity substates following DMT and placebo administration by capitalizing on neuroimaging data from a pharmacological study conducted in 14 healthy volunteers (Timmermann et al., 2023b). We applied community detection algorithms to the group-averaged subtraction matrix (DMT minus placebo) of time-resolved activity similarity (**Figure 1A-E**), to identify temporal modules of homogenous brain activity – referred to henceforth as brain activity substates – differentiating the DMT from the placebo condition.

We employed a data-driven strategy to determine the optimal Pearson’s correlation threshold to apply to the group-averaged subtraction matrix of time-resolved activity similarity, by querying for thresholds placed towards the right-end of the distribution, ranging from 0.20-0.30 (***Supplementary Figure S2A***). This approach aimed to enhance: (*i*) the identification of a maximal number of distinct brain activity substates differentiating the DMT and placebo conditions; and (ii) the discernment of brain activity substates that exhibit differential occupancy during the pre– and post-injection periods (***Supplementary Figure S2B***). A Pearson’s correlation coefficient of 0.25 was identified as the ideal threshold (five substates for a pre-post-injection separability index of *X*^2^ = 31.1), which was applied to derive five brain activity substates. Overall, overlapping brain activity substates were identified when using alternative thresholds of 0.24 (four substates for a pre-post-injection separability index of *X*^2^ = 31.6) and 0.26 (six substates for a pre-post-injection separability index of *X*^2^ = 25.2), which yielded large numbers of brain activity substates differentially occupying the pre– and post-injections periods, supporting the choice of our data-driven threshold of 0.25 (***Supplementary Figure S2C***).

State 1 and 3, although significantly more occupied during the pre-injection period, were occupied for <5% and were discarded from further analyses (**Figure 1F**). The remaining brain activity substates were occupied for substantial proportions of the scan (>10%) and their occupation significantly differed when comparing the pre-to the post-injection period (**Figure 1F**). A primary brain activity substate predominantly occurred before the injection of DMT/placebo and towards the end of the scanning session (State 2; **Figure 1E**), accounting for 34% (2.7 min) of the pre-injection scanning period and 10% (2.0 min) of the post-injection scanning period **(Figure 1F** and **Figure 2G**). It was characterized by orbitofrontal activations under both DMT and placebo (**Figure 2A-B**), while subcortical as well as medial and lateral parietal activations were identified only under DMT (**Figure 2B**). This substate was also characterized by widespread deactivation of primary sensory areas under both DMT and placebo. A second primary brain activation substate emerged shortly after the injection of DMT/placebo (State 4; **Figure 1E**) which accounted for 6% (0.5 min) of the pre-injection scanning period and 19% (3.8 min) of the post-injection scanning period **(Figure 1F** and **Figure 2G**). This substate was characterized by widespread activations of primary sensory and cingulo-opercular brain regions under both the DMT and placebo conditions (**Figure 2C-D**), while deactivations were mainly observed in prefrontal areas under both DMT and placebo and within medial parietal cortices under DMT. A third substate was identified (State 5; **Figure 1E**), which occurred immediately after the first post-injection State 4 and occupied 6% (0.5 min) of the pre-injection scanning period and 18% (3.6 min) of the post-injection scanning period **(Figure 1F** and **Figure 2G**). State 5 displayed overall high resemblance to the previous substate (**Figure 2E-F**), showing primary sensory and cingulo-opercular activations as well as prefrontal and medial parietal deactivations under DMT and placebo, yet displayed marked temporopolar activations compared to previous substates.

**Figure 2.**
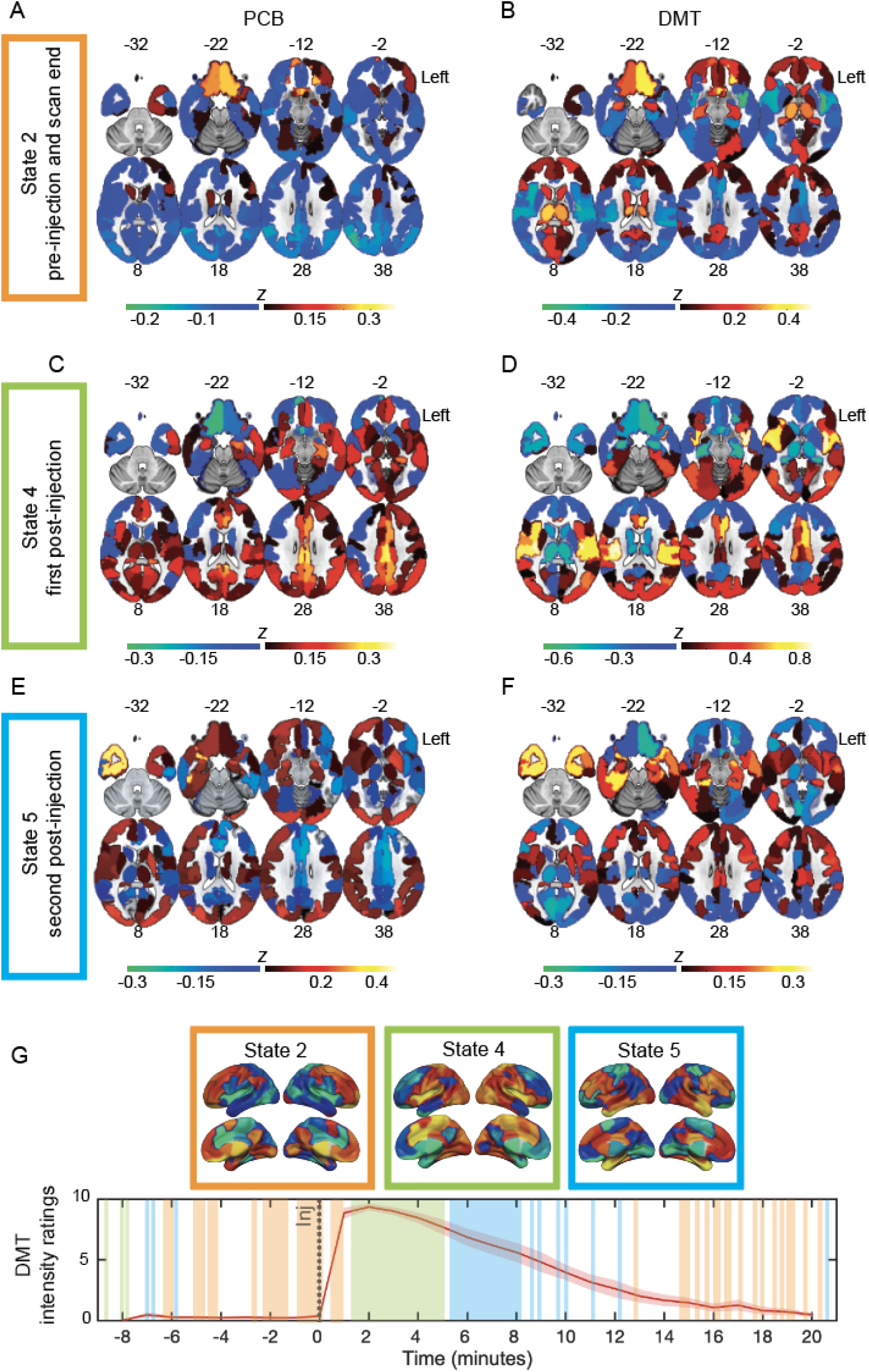
Maps of brain activity substates. Mean activation maps (z-scores) of brain substates, reflecting relative hyperactivity (warm colors) or hypoactivity (cold colors) when compared to the rest of the scan, shown for placebo (PCB, on the left) and DMT (on the right). Mean activity of State 2, which occurred before injection and towards the end of the scan, for placebo (**A**) and DMT (**B**). The duration of this substate before injection was 2.7 min, and after injection 2.0 min. Mean brain activity of State 4, occurring immediately after injection, for placebo (**C**) and DMT (**D**). The duration of this substate post-injection was 3.8 min. Mean activity of State 5, the second post-injection substate, under placebo (**E**) and DMT (**F**). The duration of this substate post-injection was 3.6 min. (**G**) Schematic representation for when brain activity substates appeared in relationship to continuous intensity ratings of the experience under DMT, assessed on a separate session in the afternoon following the acquisition of the fMRI dataset acquired in this study. Color code of rectangles framing the brain maps corresponds to the colors used to denote distinct brain activity substates in Figure 1E. The left hemisphere is shown on the right side. Inj = timepoint of injection.

### Post-injection brain substate activations and deactivations under DMT

We then used linear regression models to estimate individual activation maps for each of the brain substates and compared these maps across the DMT and placebo conditions using paired *t*-tests. Only the first post-injection brain substate (State 4) showed significant differences when comparing regional activation levels across the DMT and placebo conditions (FDR adjusted *p* < 0.05). Specifically, it displayed increased right superior temporal lobe activity under DMT, while deactivations under drug were found in the left hippocampus and in bilateral medial parietal areas overlapping with the precuneus and posterior cingulate cortices (**Figure 3A**). Crucially, mean frame-wise head displacement, a common confounder in rs-fMRI analyses, did not significantly correlate with right superior temporal lobe activity (R(*12*) = 0.00, *p* = 0.99), nor with left hippocampal and medial parietal deactivations (R(*12*) = 0.28, *p* = 0.32). Activity decreases during State 4 were not significantly predicted by the order of DMT administration (whether DMT dosing took place at the first or at the second visit; β = –0.09, *p* = 0.66, ***Supplementary Table S1***). Order of DMT administration approached significance when predicting increases in superior temporal lobe activity (β = 0.82, *p* = 0.054, ***Supplementary Table S1***), suggesting that activity increases were higher in participants receiving DMT on their first visit. To further evaluate the dynamics of activity changes under DMT, raw BOLD activity time series were averaged across voxels of State 4 displaying hyperactivity/hypoactivity under DMT. Raw activity time series were subtracted across the DMT and placebo conditions and plotted, showing sustained medial parietal and hippocampal deactivations and superior temporal lobe hyperactivations immediately after the injection of DMT (**Figure 3B**).

**Figure 3.**
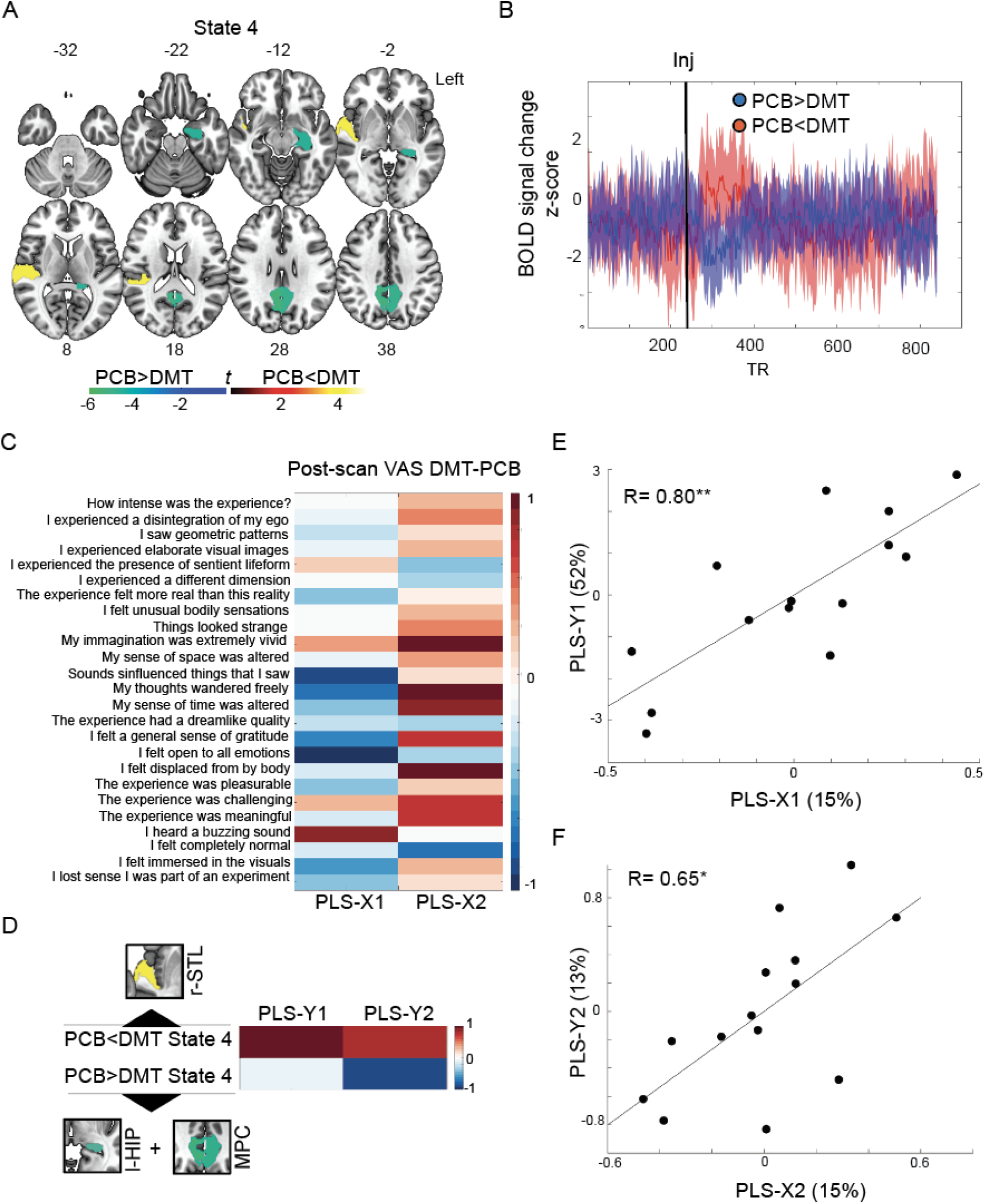
**Substate hypoactivity and hyperactivity under DMT**. **(A)** Significant hyperactivity (warm colors) and hypoactivity (cold colors) in the first post-injection substate (State 4) under DMT when compared to placebo. Findings are FDR adjusted *p* < 0.05; color bar indicates associated *t*-values. (**B**) Increased mean activity of the right superior temporal lobe (red line) and decreases in hippocampus/medial parietal activity (blue line) following DMT injection. (**C**). Loadings of questionnaire change scores on the first two partial least-square regression components. Warm colors reflect positive weights, cold colors reflect negative weights. (**D**) Loadings of right superior temporal lobe hyperactivity and left hippocampus/medial parietal hypoactivity under DMT on the first two partial least-square regression components. Warm colors reflect positive weights, cold colors reflect negative weights. Regions-of-interest used to derive averaged activity changes are shown in proximity to each contrast. (**E**) Correlation between the first dependent and independent partial least-square components. (**F**) Correlation between the second dependent and independent partial least-square components. l-HIP = left hippocampus; MPC = medial parietal cortex; r-STL = right superior temporal lobe; VAS = visual analog scale. \**p* < 0.05; \*\**p* < 0.005

We next explored the association between regional hypoactivity/hyperactivity changes under DMT and change scores of scales reflecting the quality of the psychedelic experience, which were assessed immediately after the end of the scanning session (***Supplementary Figure S4A-C).*** A partial least-square regression (**Figure 3C-F**) revealed that right superior temporal lobe hyperactivity loaded positively on a first component associated with simple and complex hallucinations, as shown by the positive loadings of questionnaires such as *“my imagination was very vivid”, “I heard a buzzing sound”*, and *“I experienced the presence of a sentient lifeform”*. In particular, the association with auditory distortions is in line with the prominent role that the superior temporal lobe plays in auditory perception (Howard et al., 2000). Hippocampal/medial parietal deactivations loaded negatively on a second component associated with alterations of essential features of consciousness and self-referential processes, indicated by positive loadings with the items *“my sense of time was altered”*, *“my sense of space was altered”,* and *“my thoughts wandered freely”*. This finding is in line with the crucial role that these regions play in the sense of time, space and self-referential processes – including the narrative self, autobiographical memories, and constructing meaning (Northoff and Bermpohl, 2004; Buckner and DiNicola, 2019), accounting for the immersive quality of the DMT experiences (Timmermann et al., 2023b). Overall, these findings suggest marked brain activity changes in the first four minutes of DMT administration, characterized by superior temporal hyperactivity, related to altered audiovisual distortions, and hippocampus/medial parietal deactivations. linked to altered sense of time, space and the self.

### Heart rate changes under DMT relate to regional brain deactivations and self-referential processes

Since autonomic bodily functions are key factors contributing to one’s sense of self (Damasio and Damasio, 2022), we proceeded to investigate changes in heart rate – a marker of sympathetic activity of the autonomic nervous system (Critchley and Harrison, 2013) – during the administration of DMT. We focused our analyses on heart rate changes occurring during the first post-injection substate, since State 4 was the only substate showing significant differences in brain activity when comparing the DMT and placebo conditions. During the first post-injection substate, heart rate was significantly increased during DMT when compared to placebo (**Figure 4A**, *t(13)* = 2.68, *p* = 0.02). Importantly, absolute mean heart rate change in State 4 did not significantly correlate with mean heart rate in State 4 under placebo (R(*12*) = –0.34, *p* = 0.24; ***Supplementary Figure S5***). A significant negative relationship would indicate that small heart rate change reflects high sympathetic tone under both the DMT and placebo condition, a factor potentially confounding our findings. Further, the order of DMT administration did not significantly predict mean heart rate increases during State 4 under DMT (β = –0.81, *p* = 0.92, ***Supplementary Table S1***). DMT-related heart rate increases during State 4 showed a significant positive correlation with State 4 hippocampus/medial parietal deactivations under DMT (**Figure 4B**, R(*12*) = 0.56, *p* = 0.04), but not with right superior temporal lobe hyperactivity under DMT (R(*12*) = 0.10, *p* = 0.72).

**Figure 4.**
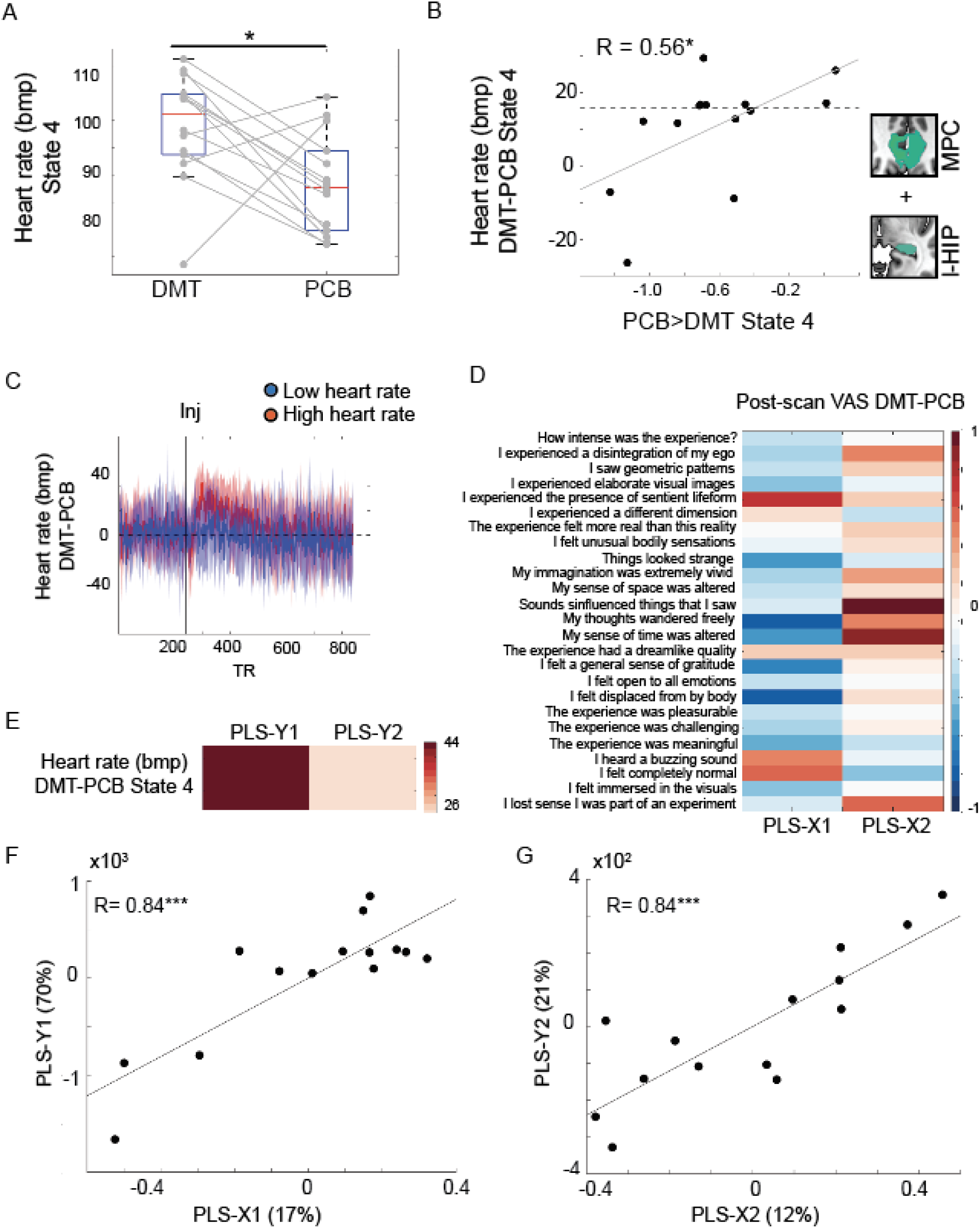
The lower heart rate under DMT, the stronger hippocampus and medial parietal deactivations. (**A**) Increased heart rate during the first post-injection activation substate (State 4) under DMT when compared to placebo. (**A**) The lower the heart rate change under the first post-injection DMT substate, the stronger hippocampus and medial parietal deactivations under the same substate. The dashed horizontal line separates participants with high heart rate during the first post-injection DMT state from participants with low heart rate based on a median split. Regions-of-interest used to derive activity decreases under DMT are shown to the right of panel **B** (l-HIP = left hippocampus; MPC = medial parietal cortex). (**C**) Continuous heart rate for participants with high heart rate (in red) and low heart rate (in blue) during the first post-injection DMT substate. Note the marked fluctuations and sustained increases in heart rate following the injection among participants with high heart rate change during State 4. (**D**). Loadings of questionnaire change scores on the first two partial least-square regression components. Warm colors reflect positive weights, cold colors reflect negative weights. (**E**) Loadings of heart rate change during the first post-injection DMT substate on the first two partial least-square regression components. Warm colors reflect positive weights, cold colors reflect negative weights. (**F**) Correlation between the first dependent and independent partial least-square components. (**G**) Correlation between the second dependent and independent partial least-square components. Bpm = beats per minute; VAS = visual analog scale. \**p* < 0.05; \*\*\**p* < 0.0005

We then used a median split to separate participants into individuals showing low heart rate increases or high heart rate increases on DMT for the duration of State 4. Continuous heart rate time series from the DMT and placebo conditions were subtracted to estimate continuous time series of heart rate change for the duration of the entire scan. Group-averaged subtracted heart rate time series were plotted separately for individuals showing low and high heart rate changes (**Figure 4C**). These plots revealed sustained heart rate increases post-DMT injection in participants showing high heart rate changes during State 4, while individuals with low heart rate changes during State 4 showed overall stable heart rate across the duration of the entire scan.

Finally, a second partial least-square regression (**Figure 4D-G**) revealed that increased heart rate changes during State 4 under DMT primarily loaded on a first component positively associated with audiovisual distortions, such as *“I heard a buzzing sound”*, which was also negatively associated with self-referential and affective functions, such as *“my thoughts wandered freely”*, *“I felt a general sense of gratitude”*, and *“I felt open to all emotions”*. These findings would suggest that participants with the strongest sympathetic activation experience more intense audiovisual distortions accompanied by a less pleasant psychedelic experience. Heart rate changes showed also positive loadings on a second component, although to a much lower degree, associated with questionnaires such as *“sounds influenced the things that I saw”* and *“I lost sense I was part of an experiment”*.

In line with previous work, our findings corroborate that the peak DMT experience is characterized by increased sympathetic output (Strassman and Qualls, 1994). Yet, lower heart rate increases are linked to a more positive psychedelic experience and stronger deactivations of brain regions related to self-referential functions.

## Discussion

To date, most efforts exploring acute brain changes during psychedelic administration have relied on metrics that capture the topology of node-to-node functional connectivity changes estimated for the duration of the entire scanning time (McCulloch et al., 2022). Yet, there is little understanding for how psychedelics may impact relative BOLD signal changes, as indexed via e.g., task-based fMRI (Smith et al., 2009). Further, there is an increased recognition that the brain displays highly dynamic properties, which underlie the emergence of complex behaviors (Calhoun et al., 2014; Cabral et al., 2017; Saggar et al., 2022). Here, we took advantage of the intrinsic dynamics of a rapidly acting psychedelic, DMT, and paired this with data-driven analytical approaches combining dynamic analyses (Kringelbach and Deco, 2020) and graph theoretical techniques (Rubinov and Sporns, 2010) to identify brain activity substates under DMT.

By comparing the DMT to the placebo condition, this analytical approach allowed us to identify sequential brain activity substates naturally unfolding under the administration of DMT. A primary brain activity substate was identified before the injection of DMT, characterized by hyperactivity of DMN areas when compared to the rest of the scan, as expected for a resting-state condition (Buckner and DiNicola, 2019). The presence of this substate at the beginning of the scanning session may reflect normal waking consciousness anchored by habitual DMN functioning, relative to brain substates and consciousness changes emerging under DMT. This DMN-based substate also occurred towards the end of the scanning session, likely reflecting the recovery from the psychedelic state back to the normal or *“default”* mode of waking consciousness (Raichle et al., 2001).

Two brain activity substates were identified under the post-DMT injection phase. The first brain substate emerged immediately after injection and lasted for several minutes. This substate was characterized by increased activity in regions subserving sensory, attentional, and interoceptive functions (Dosenbach et al., 2007; Critchley and Harrison, 2013), while deactivations were particularly prominent in medial parietal and medial temporal areas. This state is suggestive of peak DMT effects related to a state of sensory immersion and inhibition of metacognitive functions. A final, less stable brain substate was identified which temporally succeeded the previously described substate, displaying marked anterior temporal pole hyperactivity, possibly related to the attribution of semantic meaning (Younes and Gorno-Tempini, 2021). This is consistent with previous phenomenological analysis showing a recovery of higher-level qualities of cognition after the immersive peak DMT experience has subsided (Timmermann et al., 2019). Statistical comparisons of the brain activity maps between the DMT and the placebo conditions, confirmed significant changes for the first post-injection DMT brain substate, by revealing superior temporal lobe hyperactivity and medial parietal/hippocampal hypoactivity under DMT.

Our chosen statistical threshold prevented us from finding more widespread activity changes when comparing other brain substates across DMT and placebo, possibly due to the relatively small sample size. Furthermore, psychedelic experiences are highly subjective and heterogenous (Prugger et al., 2022), with factors such as the participant mindset, the cultural and social background, as well as the physical context (i.e., *“set-and-setting”*) of a session heavily influencing the perceptions, emotions, and insights elicited in different individuals (Hartogsohn, 2017; Carhart-Harris et al., 2018). This intrinsic heterogeneity may represent a challenge for the reliable identification of brain substates occurring during a psychedelic experience. Yet, medial parietal and hippocampal deactivations during peak effects may represent a signature activity pattern common to most psychedelics (Carhart-Harris et al., 2014; Tagliazucchi et al., 2014). Future work repeatedly assessing the same participant (Poldrack et al., 2015; Gratton et al., 2018; Siegel et al., 2023) across different days and dosing protocols may shed light on the inter– and intra-individual neural variability of psychedelic experiences.

When relating activity changes in brain substates following the injection of DMT to phenomenological scores acquired after the psychedelic experience, our analyses revealed that right superior temporal lobe hyperactivity correlated with visual imagery, auditory hallucinations, and the experience of entities, reflecting the emergence of unusual sensory experiences under DMT. The superior temporal lobe is a key component of the auditory cortex and plays a critical role in hearing, speech, language (Howard et al., 2000). Smaller volume in this area has been related to auditory hallucinations in schizophrenia (Barta et al., 1990) and its activity has been associated with the emergence of false auditory perceptions (Moseley et al., 2014). Overall, our findings may relate to previous reports showing increased functional connectivity of primary sensory areas, including auditory cortices, under the acute administration of DMT and other psychedelics (Tagliazucchi et al., 2016; Timmermann et al., 2023b). While hyperactivity related to the emergence of unusual sensory experiences such as audiovisual hallucinations and the experience of entities, hypoactivity in medial parietal cortices (posterior cingulate and precuneus) and the hippocampus was related to the deconstruction of essential features of consciousness and the sense of self. High doses of DMT are known for inducing *“immersive’ experiences*” (Timmermann et al., 2023b) akin to virtual reality or dreaming. Our results suggest that the immersive quality of these experiences could be underpinned by deconstructions in time, space, and the self, caused in turn, by DMT-induced hippocampal and medial parietal deactivations. These regions are key components of the DMN, a brain system which has been repeatedly associated with anchoring narrative self-referential processes (Yeshurun et al., 2021). Diminished functional integrity of the DMN, in particularly of the posterior cingulate and the precuneus, was initially shown under psilocybin (Carhart-Harris et al., 2012), and has been since confirmed under LSD (Carhart-Harris et al., 2016), ayahuasca (Palhano-Fontes et al., 2015), and DMT (Timmermann et al., 2023b). Specifically, psilocybin has been shown to reduce cerebral blood flow and BOLD activity in medial parietal areas (Carhart-Harris et al., 2012), in line with the DMT-induced midline parietal deactivations found in our study. Recent human intracranial electrophysiological studies have revealed that posteromedial cortical rhythms play a crucial role in sustaining self-referential functions as proven by both pathological seizure-based (Lyu et al., 2023) and ketamine-induced dissociations (Vesuna et al., 2020). When considering the role of hippocampal deactivations, previous rs-fMRI studies found that psychedelics decrease the amplitude of spontaneous BOLD signal fluctuations within the parahippocampal gyri and induce an uncoupling between the hippocampus/parahippocampus and other DMN regions (Carhart-Harris et al., 2014). Further, spontaneous connectivity changes within the hippocampus, parahippocampus, and medial parietal regions have been found to relate to the so-called *“ego-dissolving”* experience of psychedelics (Lebedev et al., 2015; Carhart-Harris et al., 2016). Potentially consistent with our present findings with DMT, a recent spectroscopy study found lower levels of glutamate metabolism in the hippocampus under psilocybin to be correlated with its ego-dissolving properties (Mason et al., 2020).

These findings collectively point towards the relevance of medial temporal lobe structures for the maintenance of normal waking consciousness and the ordinary sense of self that accompanies it. We found deactivation of the hippocampus to be correlated with an altered sense of time, space and self-referential processes experienced under DMT, supporting the notion that the hippocampus plays a crucial role in self-related functioning and consciousness more broadly (Northoff and Bermpohl, 2004; Carhart-Harris and Friston, 2010; Buckner and DiNicola, 2019). Further studies are required to test the relationship between self-referential functions and medial parietal/hippocampal activity patterns naturally unfolding under psychedelic administration.

Corroborating previous work on DMT and related psychedelics (Strassman and Qualls, 1994), our study revealed that the early-to-peak phase of the DMT experience is accompanied by increased heart rate, a common indicator of heightened sympathetic tone (Critchley and Harrison, 2013). Yet, participants with the highest heart rate increases under DMT showed weaker hippocampal/medial parietal deactivations. Crucially, mean heart rate assessed during the peak DMT effects was further used to differentiate participants showing either sustained increased or normal heart rate after DMT injection. Our findings are in line with human intracranial electrophysiological studies showing that medial parietal areas, including the anterior precuneus, are causally involved in the processing of the bodily sense of self (Lyu et al., 2023).

Successful regulation of the sympathetic nervous system may be mechanistically linked to midline parietal and temporal deactivation patterns (Beissner et al., 2013) underlying altered bodily experiences commonly induced by psychedelics, including ego dissolution (Carhart-Harris et al., 2012; Muthukumaraswamy et al., 2013; Lebedev et al., 2015) and self-dissociation (Lyu et al., 2023). Furthermore, those with the more modest heart rate increases were those who rated highest for altered self-referential and affective functions. Overall, our findings suggests that sympathetic regulation is a fundamental mechanism relevant to the emergence of neural activity patterns underlying self-referential processes. Intuitively, our findings can be linked to cognitive frameworks commonly used in the psychedelic field, often indicated by terms such as *“acceptance”*, *“letting-go”*, or *“surrender”*, which have been shown to be key mediators of improved well-being following assisted psychedelic therapy (Roseman et al., 2018; Wolff et al., 2020; Yaden and Griffiths, 2021). Furthermore, these findings are consistent with a complementary analysis of continuous self-report data acquired in the framework of this study showing how autonomic system fluctuations are related to acute peak experiences and increases in well-being following DMT administration compared to placebo (Bonnelle et al., 2024). The parasympathetic and sympathetic systems play a crucial role in shaping human emotions (Pasquini et al., 2022) and social behavior (Critchley and Harrison, 2013) through direct and indirect neural pathways underlying interoceptive processes and the homeostatic control of internal bodily states (Critchley and Harrison, 2013; Damasio and Damasio, 2022). We hope that our findings will stimulate future clinical studies elucidating the role of autonomic changes in predicting outcomes following psychedelic therapy (Bonnelle et al., 2024), as well as multimodal neuroscience studies exploring the dynamic integration of bodily signals within specific brain circuits under acute psychedelic administration.

## Ending Section

### Contributions

L.P. and C.T. conceived the study. R.L.C-H, L.R., H.K. and C.T. acquired the data. L.P. designed and developed the methodologies. L.P., C.T., and A.J.S. analyzed the data. L.P. and C.T. wrote the paper. L.P., A.G., C.L.G., C.T., R.L.C-H. interpreted the findings. All authors revised the paper and provided critical feedback.

### Funding

This work was supported by the following agencies: L.P.: K99-AG065457 (NIA), R00AG065457 (NIA), Feodor Lynen Research Fellowship from the Humboldt Foundation, and philanthropic support from David Dolby and the Dolby family.

### Code and Data Availability

Code for deriving brain activity substates is available on GitHub (https://github.com/lollopasquini/DMTsubstates). An unthresholded *t*-map for State 4 comparing DMT to placebo is available on NeuroVault (Gorgolewski et al., 2015) (https://identifiers.org/neurovault.image:858860). Nodal activity time series of the analyzed participants are available on GitHub (https://github.com/timmer500/DMT_Imaging).

