## Supplement for "Dynamic medial parietal and hippocampal deactivations under DMT relate to sympathetic output and altered sense of time, space, and the self"

We next explored the association between regional hypoactivity/hyperactivity changes under DMT and change scores of scales reflecting the quality of the psychedelic experience, which were assessed immediately after the end of the scanning session (***Supplementary Figure S4A-C).*** A partial least-square regression (**Figure 3C-F**) revealed that right superior temporal lobe hyperactivity loaded positively on a first component positively associated with visual and auditory hallucinations, as shown by the positive loadings of questionnaires such as *“I heard a buzzing sound”*, *“I experience the presence of sentient lifeforms”*, or *“my imagination was very vivid”*. In particular the association with auditory distortions is in line with the prominent role that the superior temporal lobe plays in auditory perception (Howard et al., 2000). Hippocampal/medial parietal deactivations loaded negatively on a second component associated with self-referential processes, as indicated by positive loadings of questionnaires such as *“my thoughts wandered freely”*, *“my sense of time was altered”*, or *“my sense of space was altered”*. This finding is in line with the crucial role that these regions play in self-referential processes, including the narrative self, autobiographical memories, and constructing meaning (Northoff and Bermpohl, 2004; Buckner and DiNicola, 2019). Overall, these findings suggest marked brain activity changes in the first four minutes of DMT administration, characterized by superior temporal hyperactivity, related to altered acoustic distortions, and hippocampus/medial parietal deactivations. linked to altered self-referential processes.

**Ending Section:**

***Conflict of interest:*** LP is a scientific advisor for AWEAR LLC.

***Contributions:*** L.P. and C.T. conceived the study. R.L.C-H, L.R., H.K. and C.T. acquired the data. L.P. designed and developed the methodologies. L.P., C.T., and A.J.S. analyzed the data. L.P. and C.T. wrote the paper. L.P., A.G., C.L.G., C.T., R.L.C-H. interpreted the findings. All authors revised the paper and provided critical feedback.

***Funding*:** This work was supported by the following agencies: L.P.: K99-AG065457 (NIA), R00AG065457 (NIA), Feodor Lynen Research Fellowship from the Humboldt Foundation, and philanthropic support from David Dolby and the Dolby family.

***Code and Data Availability****:* Code for deriving brain activity substates is available on GitHub (<https://github.com/lollopasquini>/DMTsubstates). An unthresholded *t*-map for State 4 comparing DMT to placebo is available on NeuroVault (Gorgolewski et al., 2015) (<https://identifiers.org/neurovault.image:858860>). Nodal activity time series of the analyzed participants are available on GitHub (<https://github.com/timmer500/DMT_Imaging>).

**Figures, Legends, and Tables:**

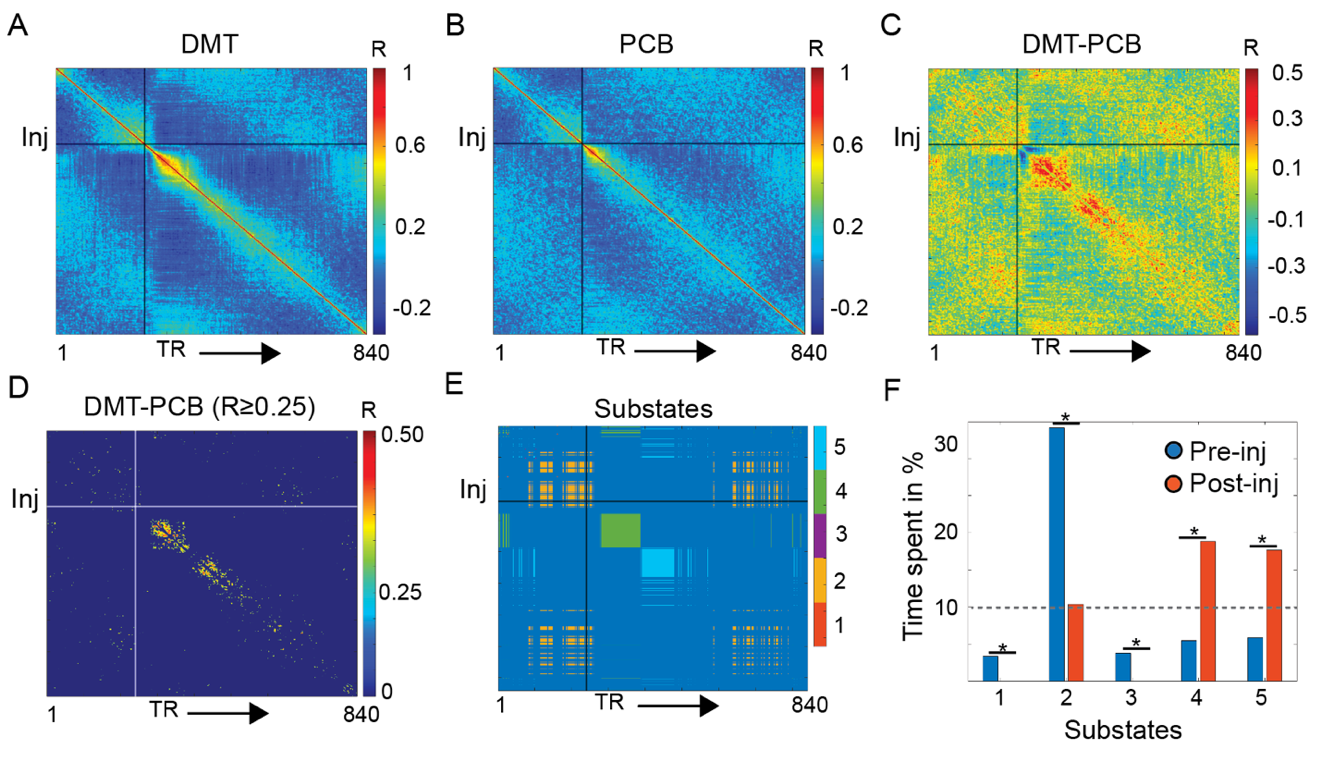

**Figure 1. Analysis pipeline and metrics.** Group-mean continuous brain activity similarity matrix, reflecting the homogeneity of brain activity in time during the DMT (**A**) and the placebo (PCB; **B**) conditions. (**C**) Mean subtraction matrix reflecting how continuous brain activity similarity varies across the DMT and placebo conditions. Warm colors reflect higher similarity of brain activity during the DMT condition. (**D**) The subtraction matrix was thresholded for similarity values maximizing the identification of separate brain activity substates occurring either during the pre-injection or post-injection periods (Pearson’s correlation value R ≥ 0.25) using an unsupervised community detection algorithm. This community detection algorithm identified five brain activity substates differentiating the DMT from the placebo condition (**E**). (**F**) These five substates were occupied at significant higher rates either during the pre- or post-injection periods, but only three out of these five substates were occupied for more that 10% of the duration of the pre- or post-injection scanning time. Following analyses focused on these three activation substates: State 2, 4, and 5. DMT = N,N-Dimethyltryptamine; Inj = timepoint of injection, indicated by continuous vertical and horizontal lines; TR = fMRI volume acquired at each repetition time. **p* < 0.05

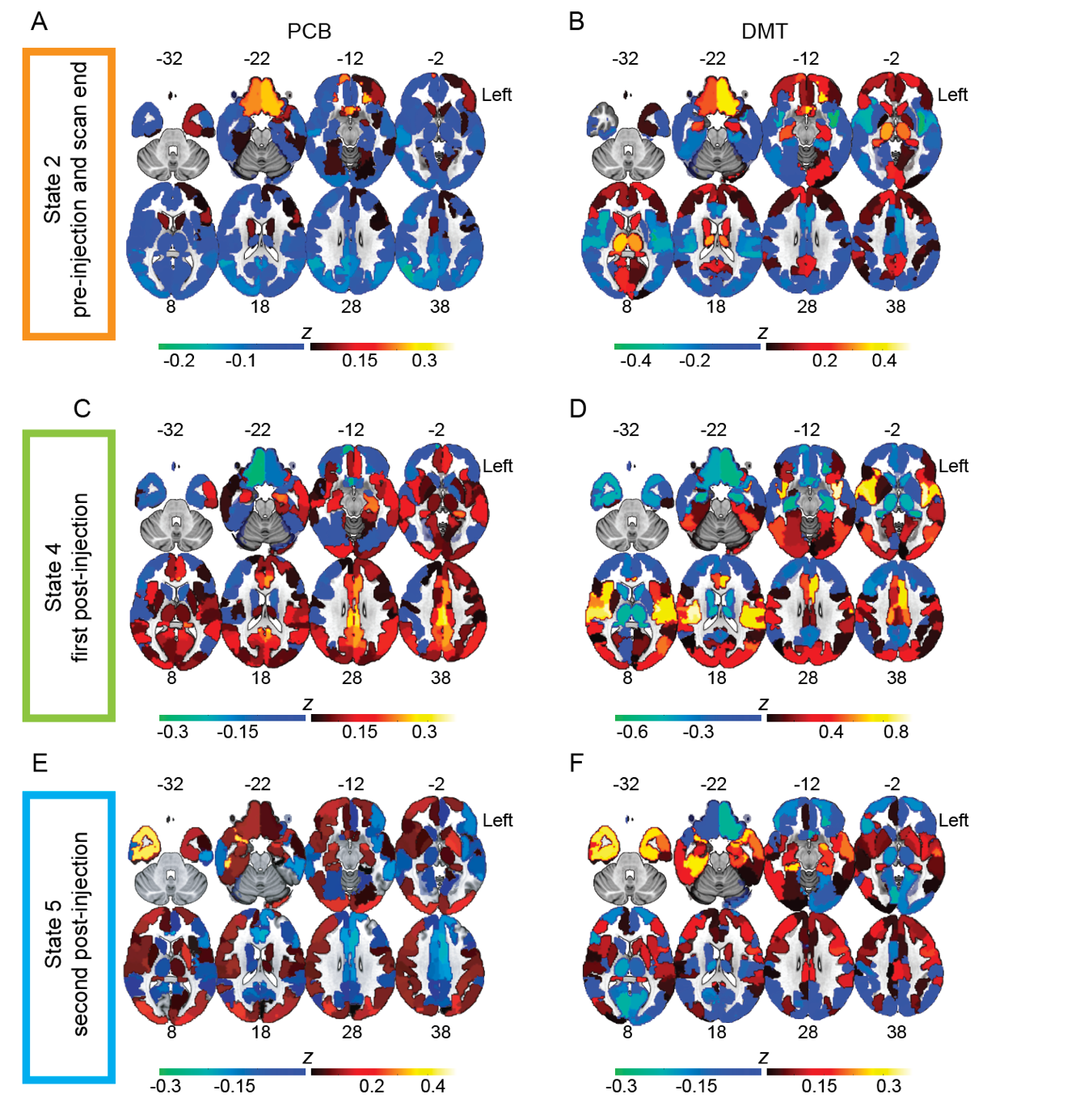

**Figure 2. Maps of brain activity substates.** Mean activation maps (z-scores) of brain substates, reflecting relative hyperactivity (warm colors) or hypoactivity (cold colors) when compared to the rest of the scan, shown once for placebo (PCB, on the left) and once for DMT (on the right). Mean activity of State 2, which occurred before injection and towards the end of the scan, once for placebo (**A**) and once for DMT (**B**). The duration of this substate before injection was 2.7 min, after injection 2.0 min. Mean brain activity of State 4, occurring immediately after injection, once for placebo (**C**) and once for DMT (**D**). The duration of this substate post-injection was 3.8 min. Mean activity of State 5, the second post-injection substate, once under placebo (**E**) and once under DMT (**F**). The duration of this substate post-injection was 3.6 min. Color code of rectangles framing the brain maps corresponds to the colors used to denote distinct brain activity substates in **Figure 1E**. The left hemisphere is shown on the right side.

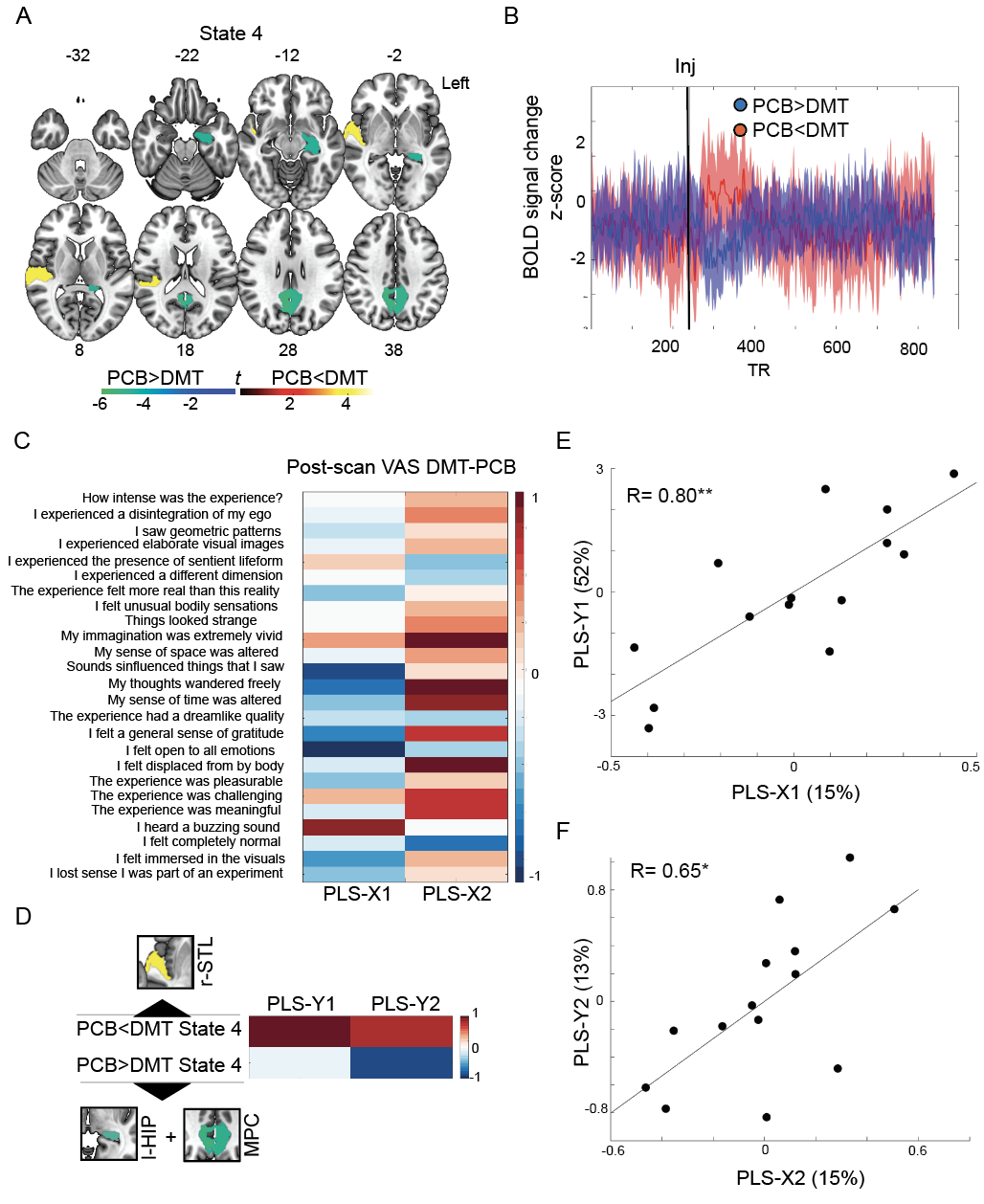

**Figure 3. Substate hypoactivity and hyperactivity under DMT.** (**A**) Significant hyperactivity (warm colors) and hypoactivity (cold colors) in the first post-injection substate (State 4) under DMT when compared to placebo. Findings are FDR adjusted *p* < 0.05; color bar indicates associated *t*-values. (**B**) Increased mean activity of the right superior temporal lobe (red line) and decreases in hippocampus/medial parietal activity (blue line) following DMT injection. (**C**). Loadings of questionnaire change scores on the first two partial least-square regression components. Warm colors reflect positive weights, cold colors reflect negative weights. (**D**) Loadings of right superior temporal lobe hyperactivity and left hippocampus/medial parietal hypoactivity under DMT on the first two partial least-square regression components. Warm colors reflect positive weights, cold colors reflect negative weights. Regions-of-interest used to derive averaged activity changes are shown in proximity to each contrast. (**E**) Correlation between the first dependent and independent partial least-square components. (**F**) Correlation between the second dependent and independent partial least-square components. l-HIP = left hippocampus; MPC = medial parietal cortex; r-STL = right superior temporal lobe; VAS = visual analog scale. **p* < 0.05; ***p* < 0.005

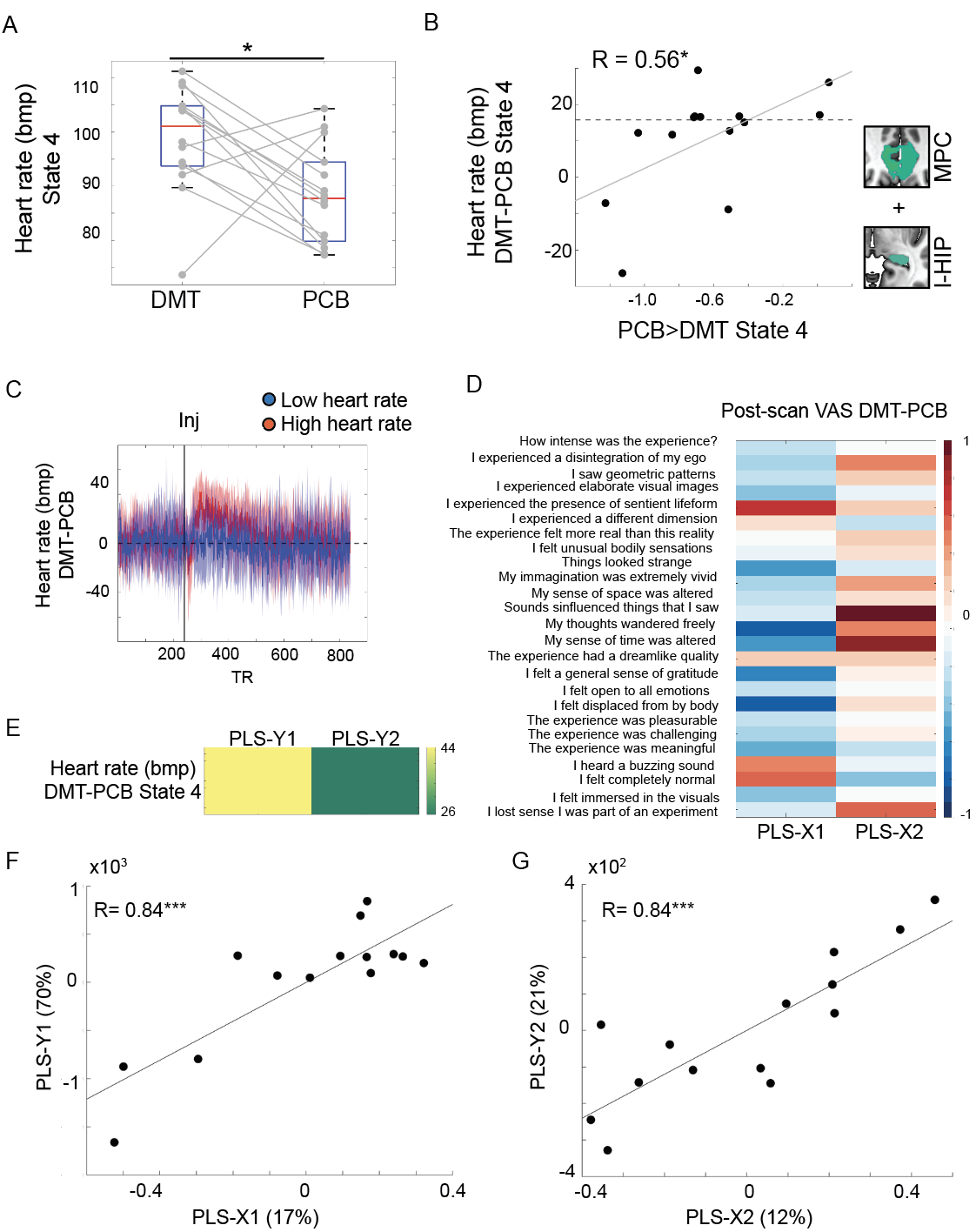

**Figure 4. The lower heart rate under DMT, the stronger hippocampus and medial parietal deactivations.** (**A**) Increased heart rate during the first post-injection activation substate (State 4) under DMT when compared to placebo. (**A**) The lower the heart rate change under the first post-injection DMT substate, the stronger hippocampus and medial parietal deactivations under the same substate. The dashed horizontal line separates participants with high heart rate during the first post-injection DMT state from participants with low heart rate based on a median split. Regions-of-interest used to derive activity decreases under DMT are shown to the right of panel **B** (l-HIP = left hippocampus; MPC = medial parietal cortex). (**C**) Continuous heart rate for participants with high heart rate (in red) and low heart rate (in blue) during the first post-injection DMT substate. Note the marked fluctuations and sustained increases in heart rate following the injection among participants with high heart rate change during State 4. (**D**). Loadings of questionnaire change scores on the first two partial least-square regression components. Warm colors reflect positive weights, cold colors reflect negative weights. (**E**) Loadings of heart rate change during the first post-injection DMT substate on the first two partial least-square regression components. Warm colors reflect positive weights, cold colors reflect negative weights. (**F**) Correlation between the first dependent and independent partial least-square components. (**G**) Correlation between the second dependent and independent partial least-square components. Bpm = beats per minute; VAS = visual analog scale. **p* < 0.05; ****p* < 0.0005
